# Moulting in Pancrustacea is characterised by both deeply conserved and recently evolved gene modules

**DOI:** 10.64898/2026.01.15.699689

**Authors:** Kenneth Kim, Giulia Campli, Sagane Joye, Ariel D. Chipman, Robert M. Waterhouse, Marc Robinson-Rechavi

## Abstract

Arthropods such as insects and crustaceans, which together form the monophyletic group Pancrustacea, possess a rigid chitinous exoskeleton that must be periodically shed through moulting to allow growth and morphological change. Although moulting is a deeply conserved developmental process across Arthropoda, our understanding of its molecular mechanisms is still largely derived from insect model species. Lineage-specific innovations and losses of moulting related genes raise fundamental questions on the extent of its conservation outside non-insect arthropods. Here, we investigate the evolutionary conservation of moulting gene expression across five representative pancrustacean species using publicly available transcriptomic datasets. Changes in gene expression during moulting are characterized by both deeply conserved and lineage-specific gene modules. Temporal gene expression analyses reveal that these lineage-specific signatures are not uniformly distributed across the moulting process: the middle transitional phase is more lineage-specific, thereby exhibiting an inverse hourglass pattern. This is likely due to life-history specific processes, development of the cuticle and specialized structures of the exoskeleton. Overall, this study provides evidence for both the evolutionary conservation and divergence of this key post-embryonic developmental process and highlights the modular architecture of the moulting programme.

## Introduction

The phylum Arthropoda includes a vast array of extant and extinct taxa, such as insects, crustaceans, arachnids, trilobites, or sea scorpions. These organisms share defining features: segmented bodies, paired jointed appendages, and a chitin-based exoskeleton, often reinforced with proteins or minerals (Chapman et al., 2013a; Daley et al., 2018; Giribet & Edgecombe, 2019). The modular organization of their segmented body plan, combined with a protective exoskeleton, has enabled arthropods to thrive in an extraordinary range of ecological niches. However, their rigid exoskeleton also hinders their ability to grow and must be periodically shed in a process called moulting, which not only enables growth but also facilitates morphological changes during development (Belles, 2020; Chapman et al., 2013a; Truman, 2019; Truman & Riddiford, 2019).

The moulting process is governed by hormonal regulation, primarily through ecdysteroids and sesquiterpenoids (Campli et al., 2024; Cheong et al., 2015; Qu et al., 2018; Truman & Riddiford, 2019). Hormones are synthesized in secretory organs in response to signals from neuropeptides. Once released into the haemolymph, these hormones reach target tissues, where they trigger transcription factors responsible for various moulting pathways (Truman, 2019). As hormone levels decline, neurosecretions activate the "ecdysis motor", a behavioural sequence that leads to the final stages of moulting and exoskeleton hardening (Roer et al., 2015; White & Ewer, 2014; Zieger et al., 2021).

Moulting is coordinated by the increase and decrease of ecdysteroid hormone or its active form 20-hydroxyecdysone (20E) (Truman & Riddiford, 2019) and occurs in distinct moult stages each characterized by different morphological and behavioural changes (Drach & Tchernigovtzeff, 1967; Gorissen & Sandeman, 2022; Kim et al., 2024; Mykles, 2011). During the pre-moult phase enzymes digest the inner layers of the old cuticle, allowing the exoskeleton to detach from the epidermal cells a process known as apolysis (Roer et al., 2015; X. Zhang et al., 2021). Moulting *sensu stricto*, or ecdysis, involves the rupturing of the old exoskeleton due to body movements and increased haemolymph pressure, enabling the animal to shed the exuvia (Daley & Drage, 2016). In the post-moult phase, the new exoskeleton hardens through processes like sclerotization, mineralization, and pigmentation, which collectively transform the soft cuticle into a mature and hardened exoskeleton.

Most of our knowledge of the molecular and biochemical mechanisms underlying moulting has been derived from a small number of insect model species, such as *Tribolium castaneum*, *Bombyx mori*, or *Drosophila melanogaster* (Campli et al., 2024; Truman & Riddiford, 2019). As a consequence, molecular insights into moulting largely stem from developmental contexts associated with insect metamorphosis, despite moulting itself being a shared and ancient process across Arthropoda and our understanding of moulting beyond Insecta remains limited (Campli et al., 2024). The increasing availability of genomic resources and sequencing data from a broader range of arthropods, driven in part by large-scale projects such as the i5k initiative (i5K Consortium, 2013) and the European Reference Genome Atlas (ERGA) (Mazzoni et al., 2023) has enabled broader testing of the involvement of genes in moulting that were first identified in insect model species (Campli et al., 2024). These genomic analyses have confirmed the presence of an conserved core set of genes involved in moulting, found across various lineages (Campli et al., 2024, 2025). While a conserved genetic backbone for moulting exists, lineage-specific modifications are evident such as the incomplete ecdysteroid biosynthetic pathway in barnacles (Dermauw et al., 2020) and the missing *Shd* (*cyp314a1*) gene in several fungus-farming ant species (Dermauw et al., 2020). Additional examples of losses include *neverland* in Varroa mites (Cabrera et al., 2015; Campli et al., 2025; Techer et al., 2019) and in beetles (Campli et al., 2025; Perry & Robin, 2025), *phantom (phm)* in chelicerates (Schumann et al., 2018), and *juvenile hormone acid methyltransferase* (*JHAMT*) gene in millipedes (So et al., 2022). Various genes also appear to be restricted to certain clades such as *Nobo* which is only found in Diptera and Lepidoptera (Enya et al., 2015; Koiwai et al., 2020), or *spok* and *Cyp6t3* found only in Drosophilidae (Ono et al., 2006; Ou et al., 2011; Sztal et al., 2007). Collectively, these patterns show that even key genes regulating moulting are not all equally conserved. The conservation of the broader set of genes involved in moulting is not well known. This is partly due to the candidate gene approach commonly used in functional studies, which prioritizes known genes from model organisms and thus leads to a systematic bias towards studying conserved genes. As most genes function within biological pathways or co-regulated modules (Brawand et al., 2011; Harrison et al., 2012; Keidar Haran & Keren, 2022; Oleksiak et al., 2002), investigating the expression patterns of genes during moulting is essential to understand the conservation and evolution of moulting processes across Arthropoda.

Phylogenetic analysis suggests that the hexapod clade, which includes insects, represents an ancient divergence within Pancrustacea, indicating that insects evolved from crustacean-like ancestors (Bernot et al., 2023; Rota-Stabelli et al., 2013; Schwentner et al., 2017; Wolfe et al., 2016). This diversity makes Pancrustacea an interesting group for studying the conservation and divergence of moulting mechanisms. Moulting in insects and crustaceans is orchestrated by neuropeptide-driven endocrine cascades regulated by specialized steroidogenic glands, the prothoracic gland (PG) in insects and the Y-organ (YO) in crustaceans (Cheong et al., 2015; Mykles, 2021; Mykles & Chang, 2020; Truman & Riddiford, 2019). In insects, moulting is triggered by the release of prothoracicotropic hormone (PTTH), which binds to the Torso receptor on the PG and activates the mitogen-activated protein kinase (MAPK) and mammalian target of rapamycin (mTOR) pathways, leading to the production of ecdysteroids (Truman & Riddiford, 2019). Both larval-to-larval and metamorphic moults are regulated by conserved endocrine pathways involving ecdysteroids and juvenile hormone (JH), with larval identity maintained by *chinmo* and metamorphic transitions associated with the sequential activation of *broad* and *E93* (Truman & Riddiford, 2026). In contrast, crustaceans rely on inhibitory control: moult-inhibiting hormone (MIH), secreted from the eyestalk X-organ/sinus gland complex, suppresses YO activity through a cAMP/Ca²⁺- and NO/cGMP-dependent signalling cascade (Mykles, 2011). Reduction of MIH release, either through environmental cues or experimental eyestalk ablation, activates the YO and triggers moulting. In addition, the calcium-rich marine environment has enabled crustaceans to build calcified, rigid exoskeletons that provide strong protection. In terrestrial environments, where calcium is less readily available, insects have adopted a different strategy by developing lighter, more flexible cuticles that rely on biochemical processes such as laccase-mediated cross-linking for cuticle hardening and tanning (Asano et al., 2019; Volovych et al., 2025). These differences in moulting mechanisms and exoskeletal properties offer a comparative framework for exploring the evolutionary conservation of moulting across Pancrustacea.

In this study, we investigated gene expression conservation of moulting using publicly available transcriptomic datasets across several insect and malacostracan species, the two largest monophyletic classes of pancrustaceans. We took two approaches to this question. First, in vertical comparison, we examined the gene expression of ecdysis *sensu stricto* and pupariation in flies (larval-prepupal moult) across five representative pancrustacean species (Table 1). Although the terminology for these processes varies in the literature, we specifically refer to moulting modes in this study as either metamorphic, such as larval-prepupal moult in insects, or non-metamorphic, i.e. the instar-instar moult. Of note, specific stages are not necessarily directly homologous across pancrustaceans. We examine shared gene expression patterns, and thus biological processes, during moulting in different species and life-stages. While larval–prepupal transitions in insects are often viewed through the lens of metamorphosis, larval-prepupal moults in insects are also hormonally driven moulting events characterized by a surge in 20-hydroxyecdysone (20E), a period of quiescence (non-feeding phase), cuticle synthesis and involve similar processes to post-ecdysial modifications such as cuticle sclerotization and tanning (Roer et al., 2015; White & Ewer, 2014). Despite the well-established role of 20E in triggering both moulting and larval-prepupal moult, downstream pathways and developmental programs that are potentially conserved between these two processes beyond insects remain largely unexplored. Secondly, in horizontal comparison, we examined gene expression conservation across several timepoints of moulting for *Drosophila melanogaster* and *Litopenaeus vannamei*, the only species in our dataset with time-series data (Supplementary Table 1). This allowed us to examine how gene expression dynamics shift across successive timepoints, and whether expression conservation patterns are consistent across the entire moulting process. We found that gene expression of moulting is characterized by both highly conserved and lineage-specific signatures, and that this pattern is not uniformly distributed. The mid-phase of the moulting is more lineage specific, exhibiting an inverse hourglass pattern. Our findings highlight the evolutionary conservation of moulting and provide insights into the evolution of moulting strategies in Pancrustacea.

**Figure 1.**
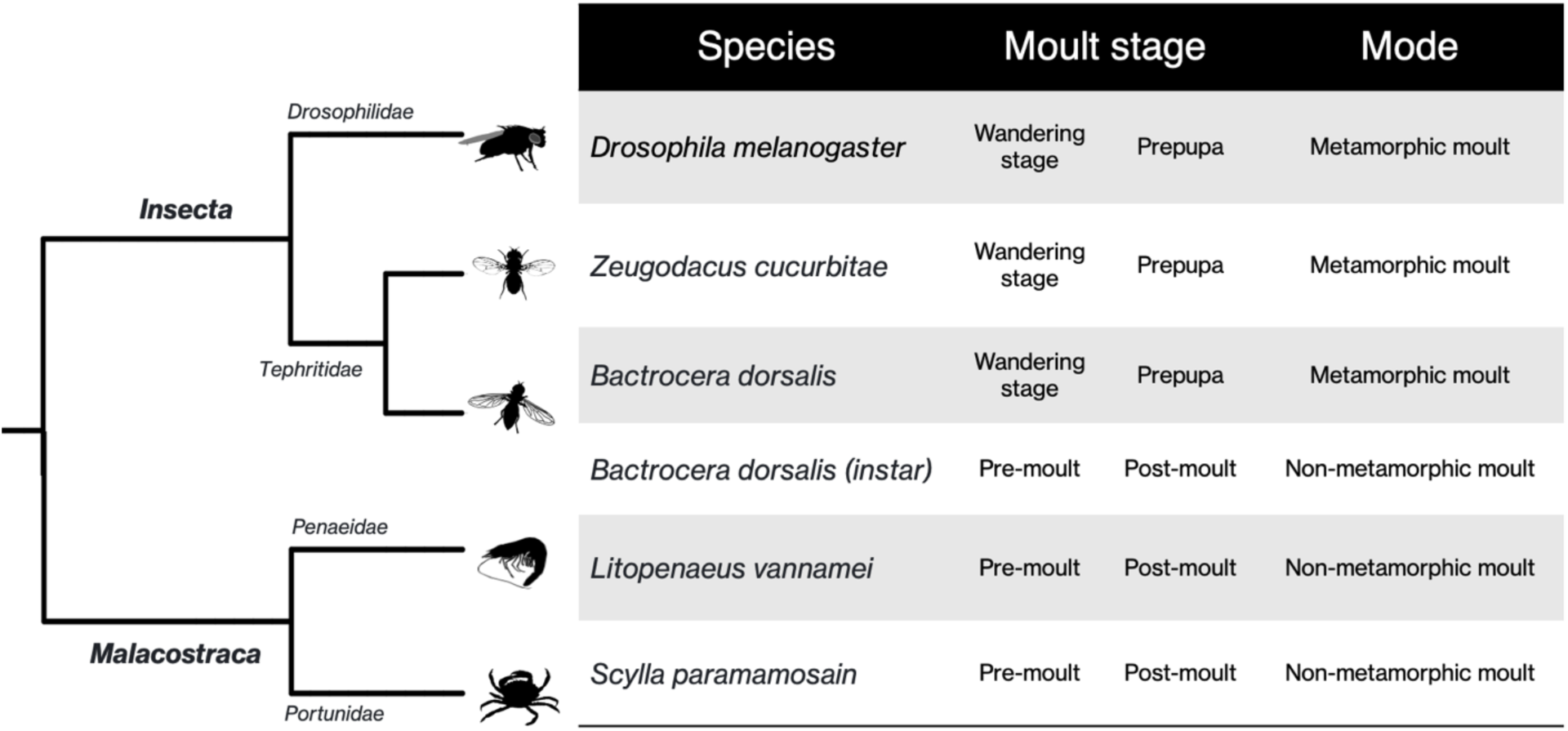
Phylogenetic tree showing the species used in this study. Additional information for each sample used is shown in the table on the right. Icon silhouettes under public domain license were retrieved from Phylopic.

**Table 1.** Summary of datasets used for gene expression analysis.

| Taxonomic group | Species | Moult stages | Source |
| --- | --- | --- | --- |
| Insecta | *Drosophila melanogaster* | Wandering stage | (Graveley et al., 2011) |
|  |  | Prepupa |  |
|  | *Zeugodacus cucurbitae* | Wandering stage | (Ward et al., 2021; Wei et al., 2020) |
|  |  | Prepupa |  |
|  | *Bactrocera dorsalis* | Wandering stage | (S. Liu et al., 2020) |
|  |  | Prepupa |  |
|  |  | Instar pre-moult |  |
|  |  | Instar post-moult |  |
| Malacostraca | *Litopenaeus vannamei* | Pre-moult | (Gao et al., 2015a) |
|  |  | Post-moult |  |
|  | *Scylla paramamosain* | Pre-moult | (L. Liu et al., 2022) |
|  |  | Post-moult |  |

## Results

### Overall gene expression patterns cluster by lineage and moult stages

To examine the gene expression pattern of moulting dataset across all samples, we performed a Principal Component Analysis (PCA) on the VST RNA-seq counts for the 5543 single-copy orthogroups identified across all species (Fig. 2). The PCA shows both clear moulting stage clustering consistent with the PCA done on individual species datasets (Supplementary Figure S7), and strong separation between insect and malacostracan lineages, indicating that lineage-specific variation represents an additional major axis of biological variation. These results indicate that the dominant patterns observed in the dataset primarily reflect biological rather than technical variation. The first 10 principal components (PCs) explained 95% of the variance in gene expression, with PC1 and PC2 alone accounting for 54%. We tested whether these PCs were significantly associated with either moult stage, subphylum, or species (Fig. 2C). The strongest correlation with moult stage was PC6 and plotting PC2 and PC6 clustered samples by moult stages. Among the genes which contribute the most to PC 6 are genes known to be involved in moulting, such as hormone receptor 3 (*Hr3)*, ecdysone induces protein 78c (*Eip78c)*, ecdysis triggering hormone receptor (*ETHR)*, and various cuticular proteins and chitinases (Supplementary Table 2). There are also uncharacterized genes, such as CG11892 which has predicted ecdysteroid-22 kinase activity, and CG12009, which has a predicted chitin binding perithophin-A domain. All samples were derived from whole-body extracts, yet despite variable involvement of different tissues in moulting, we detected a clear signal of moulting in the dataset. This likely reflects the large changes in gene expression just before and after ecdysis and pupariation.

**Figure 2.**
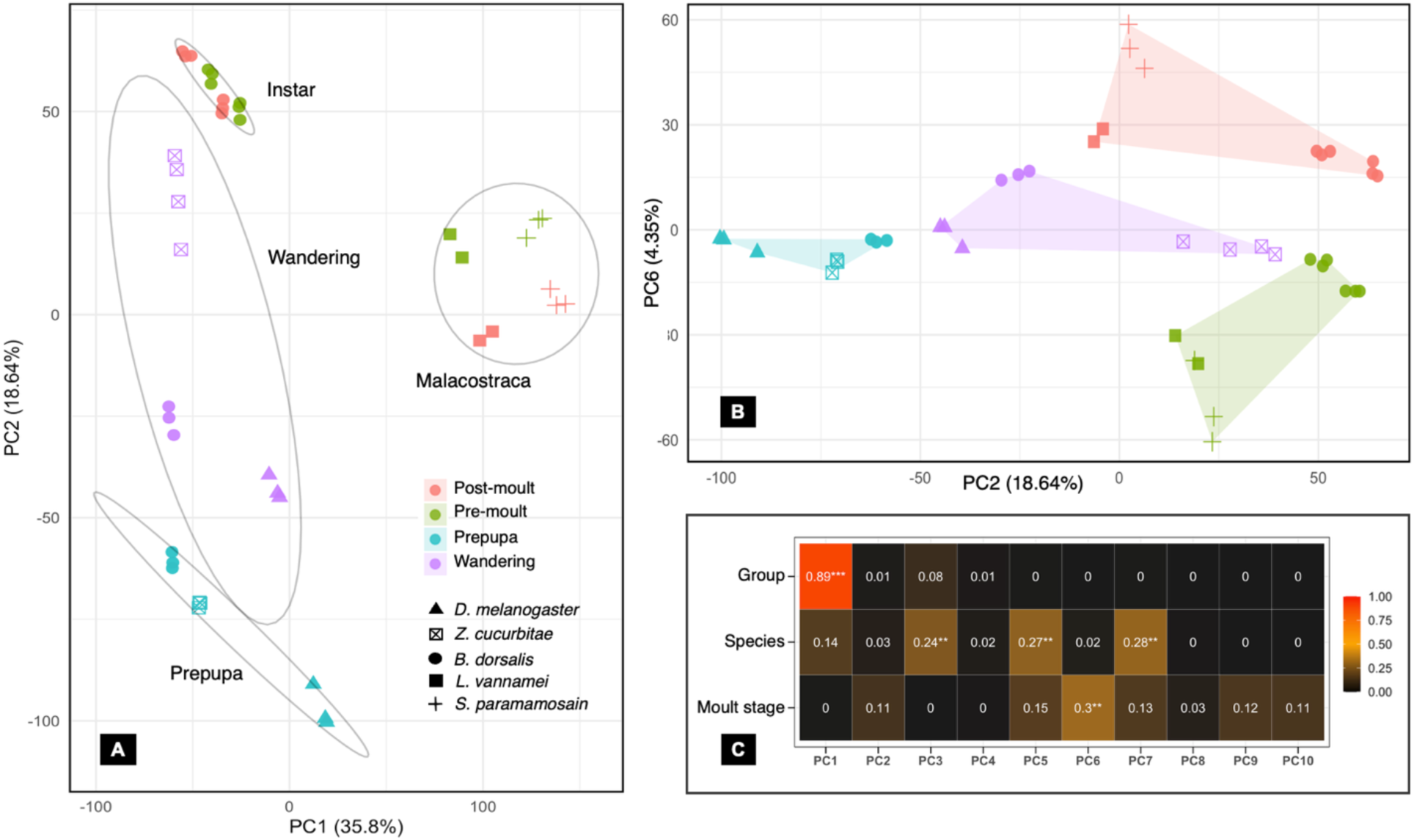
Principal Component Analysis of RNA-seq data. **A**. PC1 and PC2 cluster samples by taxonomic group and moult stage. **B**. PC2 and PC6 show clustering of samples by moult stage. Left and top portion shows prepupa and post-moult stage while the samples at the centre and bottom show wandering and pre-moult stages. **C**. Eigenvalue correlations of the principal components with respect to taxonomic group, species, and moult stage. Colour indicates the strength of the correlation and asterisks indicate level of significance after Benjamini–Hochberg (Benjamini and Hochberg 1995) correction with \**P* < 0.05, \*\**P* < 0.01, and \*\*\**P* < 0.001.

### Shared and lineage-specific differentially expressed genes in non-metamorphic moults

We first consider only the non-metamorphic moults, i.e. the instar moult of *B. dorsalis* and the two malacostracans (Figure 3, top). While the largest fraction of differentially expressed genes (DEGs) between pre- and post-moult are classified as lineage-specific (i.e. species-specific or class-specific), there are also many DEGs which are shared among pancrustaceans.

**Figure 3.**
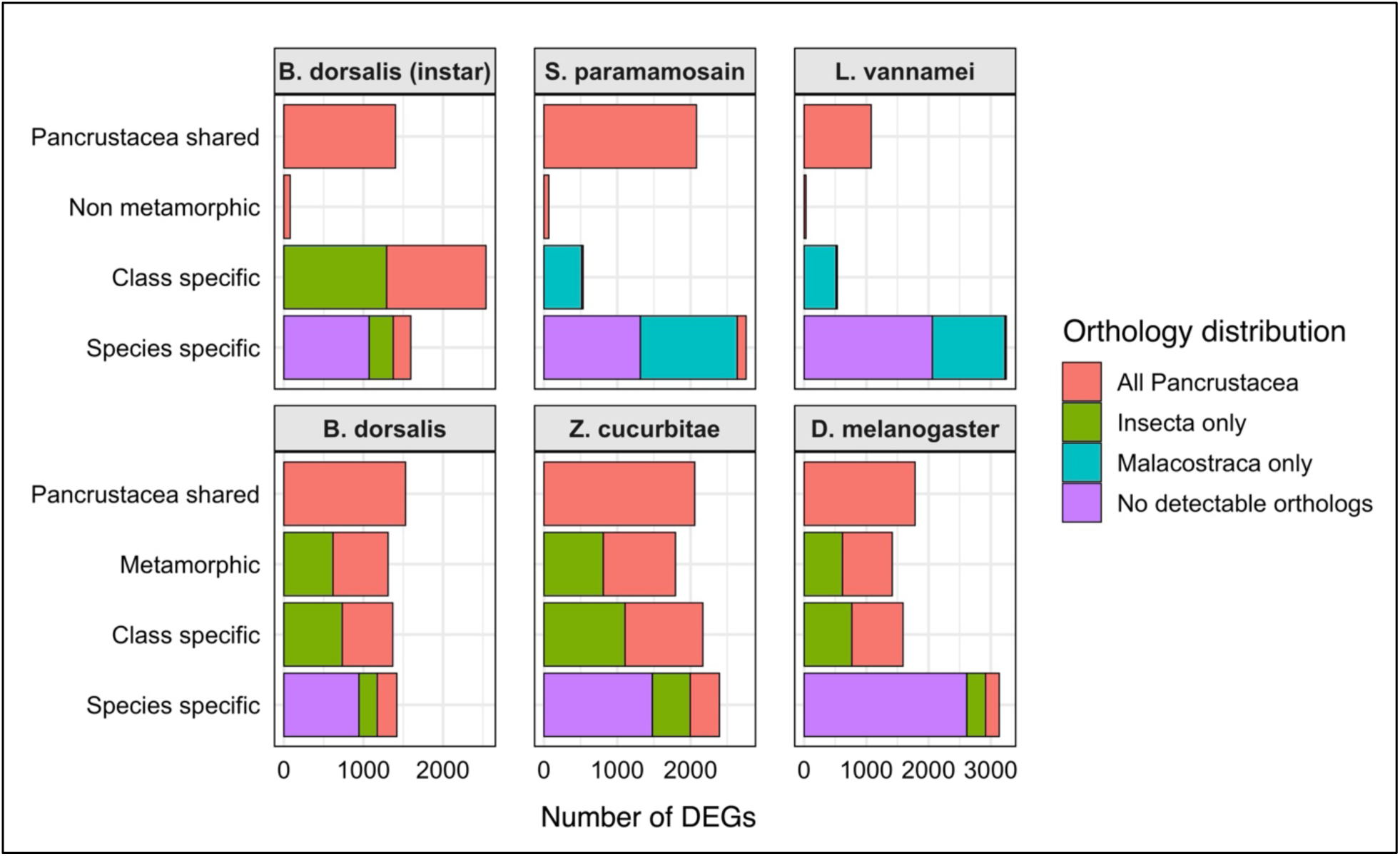
Classification of differentially expressed genes (DEGs) and their corresponding orthology distribution. Top: non metamorphic moults; bottom: metamorphic moults. "Class specific" are DEGs restricted to Malacostraca or to insects.

These shared DEGs include known members of the late genes and ecdysis neuromotor pathways, such as *knickkopf* (*Knk*), *obstructor-A* (*Obs-A*), retroactive *(Rtv), serpentine* (*serp*), *krotzkopf verkehrt* (*kkv*), and various chitin deacetylases and chitinases (Campli et al., 2024; Moussian et al., 2015; Pesch et al., 2015, 2016). In addition, many of these DEGs are involved in cuticle development, and include various esterases, ecdysteroid kinases, and structural components such as extracellular matrix proteins, gap junction elements, and other support structures (Supplementary Table 3). We also found cytochrome P450 (CYP) genes and other steroid binding proteins and transporters, some of which have been shown to be involved in ecdysone metabolism and sterol transport, such as organic anion transporter polypeptides, Niemann-Pick type C (*NPC*), δ-aminolevulinic acid synthase (*Alas*), megalin, and membrane steroid binding protein (*MSBP*) (Fujii-Taira et al., 2009; Huang et al., 2005; Nakaoka et al., 2017; Riedel et al., 2011). Many of these genes have orthologues outside Pancrustacea, indicating that their involvement in moulting might be conserved more broadly.

### Comparison of metamorphic and non-metamorphic moults

To examine whether these Pancrustacea-shared DEGs are restricted to non-metamorphic moults or are conserved more broadly across moulting modes, we extended the comparison to include metamorphic moults in insects (Figure 3). Interestingly, only a small fraction of DEGs was exclusively shared among non-metamorphic moults, whereas Pancrustacea-shared DEGs were broadly represented among DEGs in metamorphic moults. This indicates that many conserved components of the moulting programme are deployed in both moulting modes.

There were differences between insects and malacostracans in the number of species-specific compared to class-specific DEGs. Malacostracan species exhibited a higher proportion of species-specific DEGs than insects; splitting them between upregulated and downregulated genes shows similar trends (Supplementary Figure S1). This pattern likely reflects the greater divergence among species within each group. *L. vannamei* and *S. paramamosain* belong to distinct decapod groups Dendrobranchiata and Pleocyemata (Reptants), respectively, which diverged approximately 400 million years ago. They exhibit markedly different life histories and ecology with *L. vannamei* fully aquatic and *S. paramamosain* semi-terrestrial. In contrast, *B. dorsalis*, *Z*. *cucurbitae* (both Tephritidae), and *D*. *melanogaster* (Drosophilidae), diverged only around 150 million years ago (estimated using TimeTree: Kumar et al., 2022).

In addition to differences in DEG numbers, class-specific DEGs differed markedly in their orthology distribution between insects and malacostracans (Figure 3). In the two malacostracans, class-specific DEGs were predominantly composed of Malacostraca-restricted orthologous groups. In contrast, class-specific DEGs in insects included a substantial fraction of Pancrustacea-shared orthologous groups. Across all species, species-specific DEGs were strongly enriched in genes with lineage-restricted orthogroups and in genes with no detectable orthologs, indicating that transcriptional novelty during ecdysis is primarily driven by genes with lineage-restricted orthology distribution rather than by changes in expression of conserved orthologous groups.

### Gene co-expression analysis shows conserved gene expression pattern of moulting

Most genes function within biological pathways or co-expression networks, and concerted expression changes of distinct gene sets can provide valuable insights into various co-regulated gene modules involved in moulting. We performed a co-expression analysis of all 5543 orthologues across all species and identified gene clusters with similar expression patterns (Figure 4). We recovered two major clusters that exhibit consistent expression across all samples: cluster 1 consists of genes that are down-regulated during the post-moult stage (Figure 4: top panel), and cluster 2 consists of genes that are up-regulated during the post-moult stage (Figure 4: bottom panel). A total of 774 co-expressed genes (14% of all orthologues) were identified across the five species (Supplementary Table 4). When the instar moult sample from *B*. *dorsalis* was included in the analysis, the number of co-expressed genes decreased to 184 genes (3% of all orthologues) (Figure 5).

**Figure 4.**
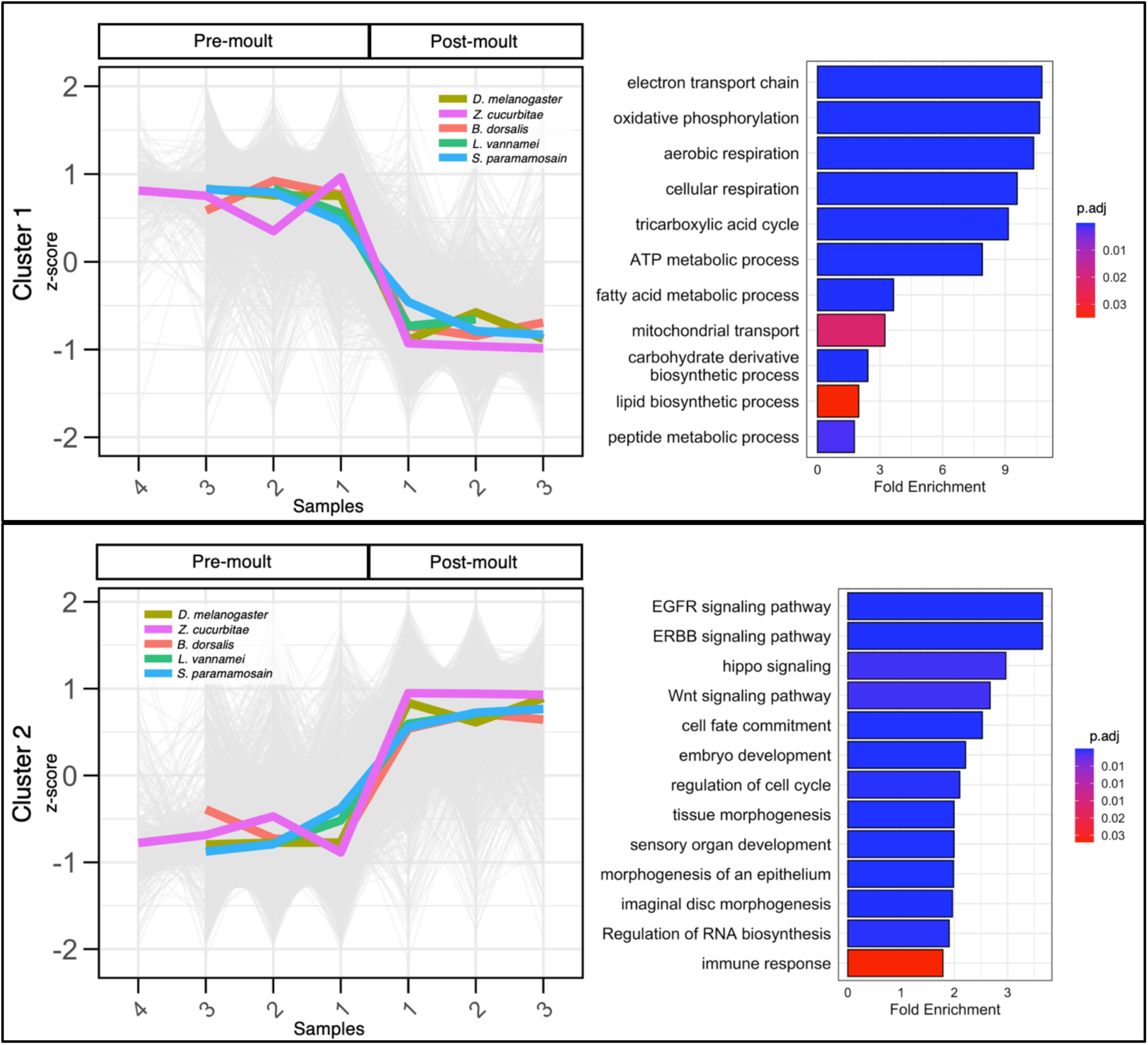
Clusters of co-expressed genes in moulting. The strength of expression in z-scores (*Y*-axis) is illustrated for the different moult stage replicates per species where coloured lines represent the expression mean while grey lines represent individual gene expression values per replicate (*X*-axis). Gene ontology (GO) enrichment plots showing representative significant terms for each cluster are shown on the right side.

**Figure 5.**
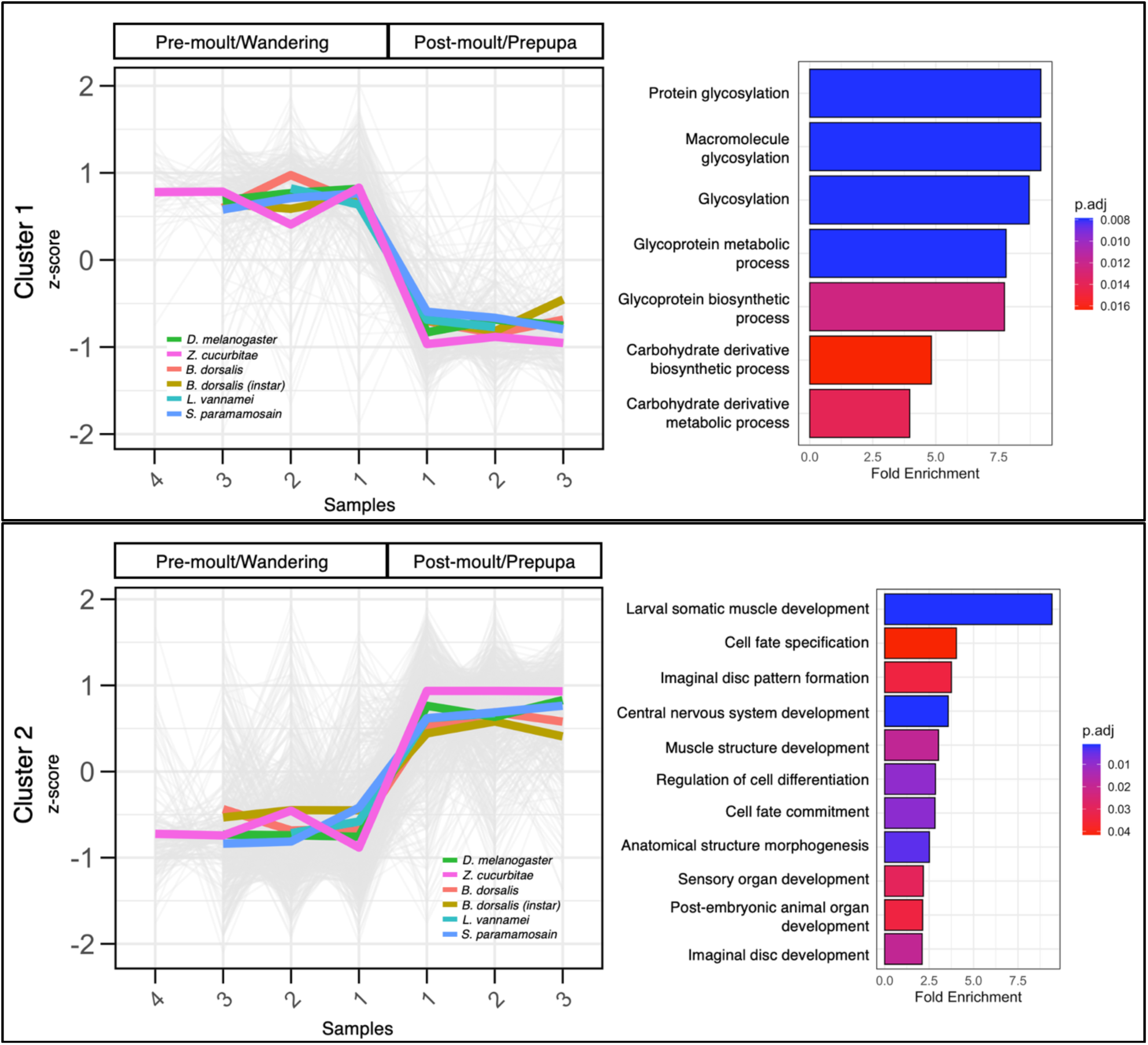
Clusters of co-expressed genes in moulting (with *B. dorsalis* instar). The strength of expression in z-scores (*Y*-axis) is illustrated for the different moult stage replicates per species where coloured lines represent the expression mean while grey lines represent individual gene expression values per replicate (*X*-axis). Gene ontology (GO) enrichment plots showing representative significant terms for each cluster are shown on the right side.

Functional analysis of the 774 co-expressed genes shows that genes in cluster 1 are enriched in terms related to core metabolic and cellular processes, such as cellular respiration and energy metabolism. Meanwhile, cluster 2 is enriched for terms related to tissue morphogenesis and developmental signalling, suggesting that the post-moult stage is characterized by the activation of pathways associated with cell proliferation, differentiation, and tissue remodelling, as expected. In addition to this we also recovered immune-related genes in cluster 2, including those known to mediate melanotic encapsulation of foreign targets, such as *zizimin-related* (*zir*), *ephexin* (*exn*), *Rho guanine nucleotide exchange factor 3* (*RhoGEF3*), and *Rho-related BTB domain-containing protein* (*RhoBTB*), and Toll and IMD signalling pathway genes including *Relish*, *supernumerary limbs* (*slmb*), *activating transcription factor-2* (*ATF-2*), *spatzle 3* (*spz3*), and *caspase* (*casp*).

The co-expressed gene sets which include the *B. dorsalis* instar-stage are not enriched for genes associated with cellular respiration or signalling pathways (Figure 5). Instead, for cluster 1 the enriched Gene Ontology terms were limited to processes such as protein glycosylation and carbohydrate metabolism. For cluster 2, we continued to recover terms related to tissue morphogenesis, sensory organ development, and muscle development. However, terms associated with key signalling pathways such as *Wnt*, *Hippo*, and *EGFR* were no longer recovered. This suggests that the metabolic and developmental requirements of instar moult in *B. dorsalis* might be different from other samples included in this analysis.

### Moulting gene expression exhibits varying patterns of conservation

Despite recovering a subset of genes that are co-expressed across all species, most genes did not show consistent co-expression patterns across species and groups. This suggests that moulting gene expression is for a large part lineage-specific, with gene expression dynamics varying between species. However, moulting as a process involves more than just the two timepoints around ecdysis. Thus, we expanded our comparison to include additional timepoints from pupariation in *D. melanogaster* and moulting in *L. vannamei*. We then examined transcriptome age patterns (TAI) across these timepoints. We found that conservation of gene expression is not evenly distributed across moulting in *L. vannamei* and larval-prepupal moult in *D. melanogaster*. For both species, TAI during the middle stages tends to be younger (i.e., more lineage-specific) showing an inverse hourglass pattern (Figures 6 and 7). However, it should be noted that while transcriptome age index (TAI) analysis is a widely used method to assess the evolutionary age of gene expression profiles across developmental stages it is also known to be sensitive to certain biases (Piasecka et al., 2013). One of the main challenges is that TAI relies on the mean expression values of genes, which can be skewed by a small number of very highly expressed genes such as housekeeping genes. These genes often have stable and abundant expression across conditions and may mask the contribution of biologically meaningful signals from genes with lower expression. As previously shown, removing or correcting for the influence of highly expressed genes can substantially alter the resulting TAI patterns, often revealing an additional layer of conservation pattern that were otherwise hidden (Piasecka et al., 2013), although this is moderated by the use of transformed expression values (J. Liu & Robinson-Rechavi, 2018). Moreover, the differences in TAI that we observe between stages are very small relative to the range of possible values. To assess whether the observed TAI patterns were driven by highly expressed but invariant genes, we repeated the analysis after filtering genes based on expression variance across stages. Specifically, we removed the lowest 10%, 25%, and 50% of genes ranked by variance and recomputed TAI. In all cases, the overall pattern was preserved, indicating that dynamically expressed genes contribute most strongly to the enrichment of younger genes during the middle stages of moulting (Supplementary Figure S8). These results suggest that the observed enrichment of evolutionarily younger genes is not simply an artefact of constitutive expression but is associated with the most dynamically expressed components of the moulting transcriptome. To further address this limitation, we also performed a temporal co-expression analysis to identify clusters of genes with similar expression patterns.

**Figure 6.**
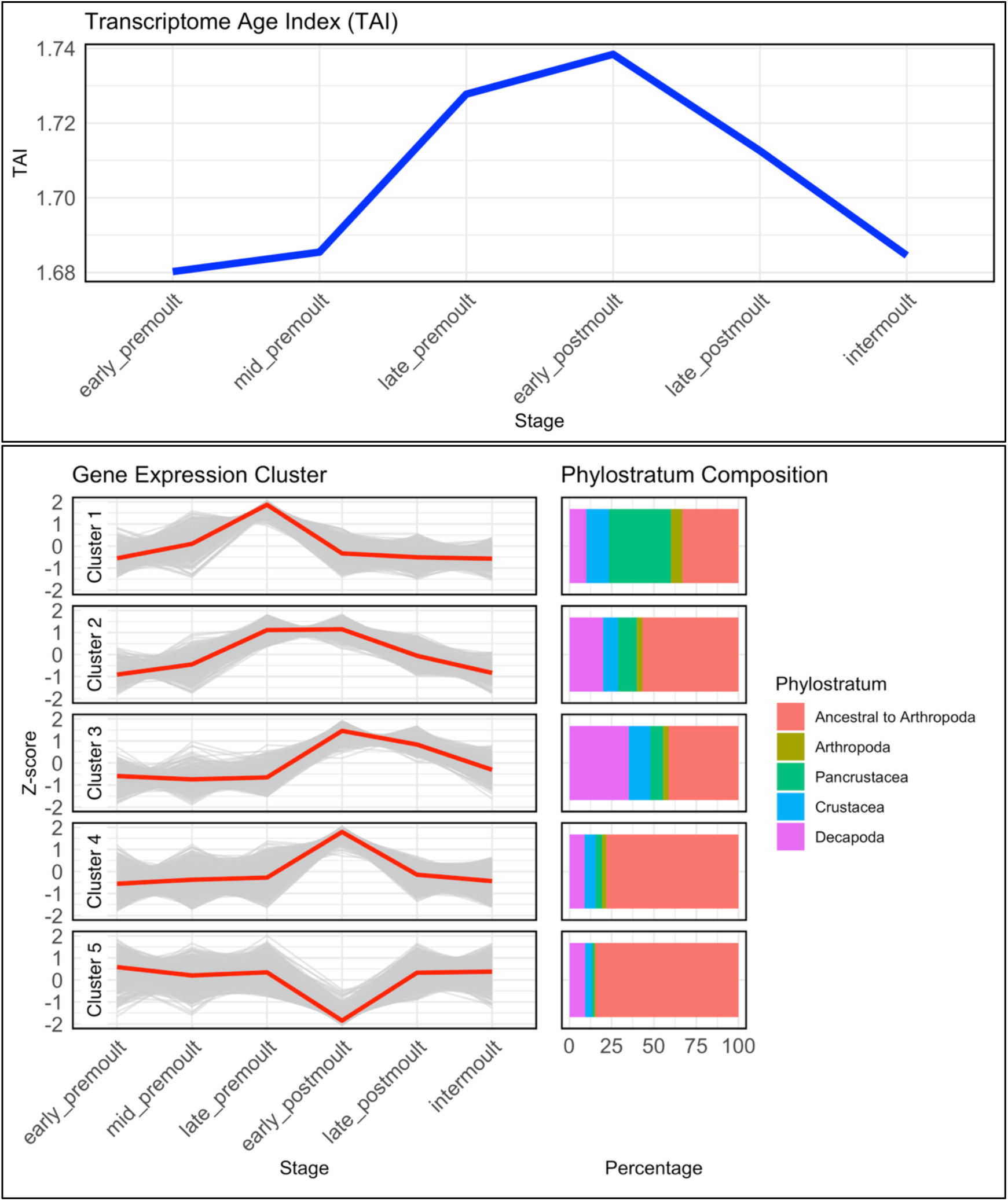
Transcriptome age index (TAI) and moulting gene expression clusters in *L. vannamei*. Top panel shows the TAI of various moult stages in *L. vannamei* (*P* < 0.0001). Bottom panel shows the gene expression clusters of *L. vannamei* moult samples and their corresponding phylostratum composition.

**Figure 7.**
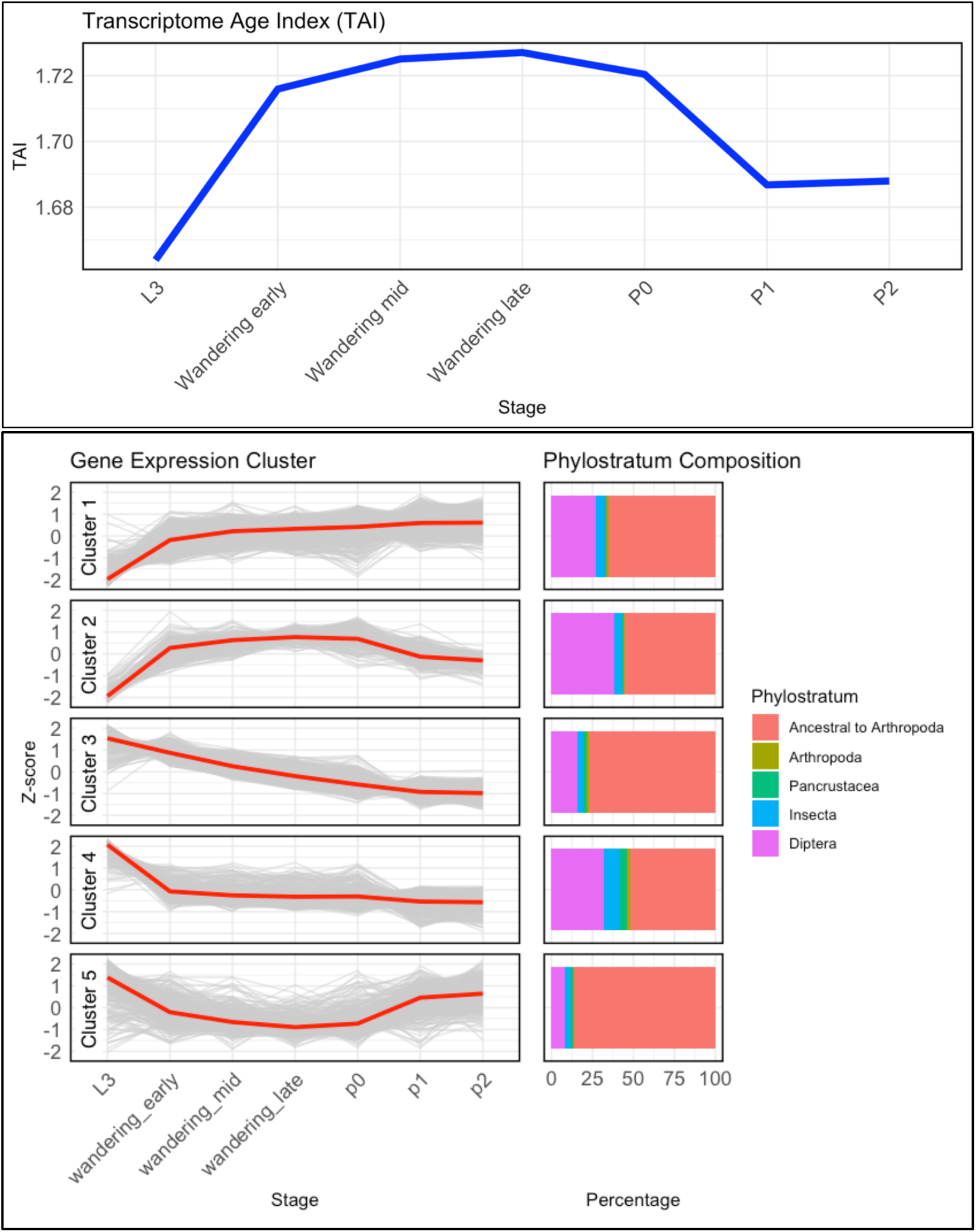
Transcriptome age index (TAI) and moulting gene expression clusters in *D. melanogaster*. Top panel shows the TAI of various moult stages in *D. melanogaster* (*P* < 0.0001). Bottom panel shows the representative gene expression clusters of *D. melanogaster* samples and their corresponding phylostratum composition.

We identified clusters of genes showing distinct trends across the moulting process (Figures 6 and 7). For each of these clusters we recovered varying patterns of conservation based on their Phylostratum composition. In *L. vannamei*, most of the gene clusters were consistent with TAI analysis. Clusters with higher expression in the middle of the moult process tend to be composed of younger genes, while those with lower expression during this phase are enriched for older, more conserved genes. For example, cluster 1, which shows peak expression in the late pre-moult stage, has a higher proportion of genes assigned to phylostrata dating back to the Pancrustacea node. In contrast, the relative contribution of more recent nodes, particularly those specific to Decapoda, increases toward the post-moult stages. On the other hand, cluster 4, which peaks in early post-moult, deviates from the overall pattern by being composed largely of older genes, suggesting a possible reactivation of conserved programmes following ecdysis. Although most gene clusters are dominated by younger phylostrata, especially during the middle of the moult cycle, it is worth noting the changes of phylostrata coming from Pancrustacea and Decapoda. This shift in phylostratum composition from a greater contribution of Pancrustacea-level genes in the late pre-moult stage to increased representation of Decapoda-level genes in the post-moult stage reflects the stage-specific recruitment of conserved and lineage-specific processes across the moulting process.

Younger gene clusters were enriched in terms involved in immune response, nervous system formation, sensory organ development, peroxidase activity, pigment binding, and cuticle development (Supplementary Figure S2). Cuticle development is enriched in clusters 1 – 3, especially in cluster 1. We confirmed this with cuticleDB (Ioannidou et al., 2014; Magkrioti et al., 2004) (Supplementary Figure S3). Genes that are predominantly expressed in cluster 3 (post-moult) are enriched in terms related to carbonate dehydratase and pigment binding, which implies the stage specific expression of genes potentially involved in calcification and pigmentation. Clusters with older phylostratum composition were enriched in terms related to extracellular matrix organization, fatty acid binding, and conserved metabolic processes such as steroid and carbohydrate metabolism, proteolysis, and lipid biosynthesis. These functional interpretations are based on Gene Ontology (GO) annotations, which can be broad and may reflect pleiotropic functions. We therefore interpret these enrichments as putative functions derived from computational annotation that require further experimental validation.

In *D. melanogaster*, we recovered ten gene expression clusters with varying temporal co-expression patterns (Figure 7; Supplementary Figure S4). The gene clusters that we recovered are largely not consistent with the TAI analysis, with mixed patterns (Figure 7). Most of the clusters exhibit a notable contribution of Diptera-specific genes, which is consistent with previous observations on the recruitment of younger genes in life stage transitions of flies (Remmel et al., 2025). Most clusters were dominated by older phylostrata and enriched in cellular and metabolic processes (Supplementary Figure S5). Cluster 1 is enriched in terms related to cilium movement, sperm flagellum assembly, cluster 2 is enriched in gamete development such as microtubule polymerization, spermatogenesis and spermatid differentiation while cluster 3 is enriched in RNA biogenesis, body morphogenesis, and cuticle development.

## Discussion

Moulting is a highly conserved developmental process across Arthropoda. While a core genetic toolkit for moulting has been identified (Campli et al., 2024, 2025), largely derived from insect model systems, comparative genomic studies suggest that this “moulting genetic backbone” has undergone lineage-specific elaborations through gene gains and losses (Campli et al., 2025). The extent to which genes involved in moulting beyond this core toolkit are conserved or lineage-specific remains poorly known. In this study, we analysed gene expression datasets from insects and malacostracans to investigate the conservation of moulting gene expression in Pancrustacea.

We identified differentially expressed genes that are conserved between distant Pancrustacea species including genes involved in ecdysteroid metabolism. In addition, we recovered genes involved in cuticle development and in post-ecdysial processes. They notably include genes involved in the structural organization of the cuticle. For example *Knk*, regulated by *Rtv*, organizes chitin into laminae and protects it from chitinase-mediated degradation, along with chitin deacetylases known to direct proper cuticle organization (Chaudhari et al., 2011, 2013; Moussian, 2013). In the extracellular matrix, *Obst-A* binds the nascent chitin fibres produced by *Kkv* and recruits the *Serp/Knk* complex to the apical plasma membrane for chitin organization (Moussian, 2013; Ostrowski et al., 2002; Pesch et al., 2015; Petkau et al., 2012). This complex is essential for aligning the chitin into an ordered structure and for protecting the developing matrix from premature degradation by chitinases. Loss of these genes results in moulting defects, chitin disorganization and lethality (Chaudhari et al., 2011, 2013; Moussian, 2013; Ostrowski et al., 2002; Pesch et al., 2015; Petkau et al., 2012).

In addition to structural organization and chitin synthesis machinery, we also found genes involved in protein trafficking and membrane transport, that are essential for cuticle turnover. The enrichment of genes related to lysosomal activity, protein trafficking, and endocytosis during ecdysis are likely due to chitin synthesis and degradation. As the old cuticle is degraded, endocytosis enables the uptake of cuticle fragments and extracellular debris, which are processed in lysosomes to recover reusable monomers, while the secretory pathways direct newly synthesized cuticular proteins, chitin synthases, and other structural components to the apical surface for assembly of the new cuticle (Moussian et al., 2007). Consistent with this, we identified core secretion pathway genes such as *ghost* (*Gho*), *Haunted* (*Hau/Sec23*), *Sec24*, and other COAT proteins, which are known to be essential for apical plasma membrane remodelling during cuticle differentiation (Moussian et al., 2007; Norum et al., 2010). The multi-ligand endocytic receptor *Megalin* (*Mgl*) in *Drosophila* illustrates another trafficking-dependent mechanism, where *Mgl*-mediated endocytosis of Yellow protein restricts pigment deposition to the distal procuticle, ensuring proper spatial organization of cuticle components (Christensen & Willnow, 1999; Moestrup & Verroust, 2001; Riedel et al., 2011). In insects, similar roles for endocytosis have been reported. In *Locusta migratoria*, clathrin-mediated endocytosis supports resource recycling during cuticle degradation, and its disruption causes moulting defects, consistent with findings in *Helicoverpa armigera* where a G protein-coupled receptor mediates 20-hydroxyecdysone uptake via clathrin-dependent endocytosis (Kang et al., 2021; Shi et al., 2022). Evidence from *Caenorhabditis elegans* shows that disruption of endocytic trafficking in epidermal cells leads to moulting arrest, defective clearance of the old cuticle, failure to internalize cargoes such as steroid hormone precursors, and disruption of plasma membrane homeostasis during new cuticle secretion (Lažetić & Fay, 2017; Yochem et al., 2015). The occurrence of such trafficking-dependent mechanisms in nematodes suggests that they represent an ancestral moulting machinery that predates Arthropoda.

Genes co-expressed during ecdysis are not limited to those directly involved in cuticle synthesis and organization. Instead, moulting also recruits evolutionarily conserved pathways supporting broader physiological functions, including metabolism, immune response, cellular differentiation, and tissue morphogenesis. Moulting is an energy-intensive process such that the caloric loss associated with shedding the exuviae is substantial (Harvey et al., 1980; Sweeney & Vannote, 1981). Additionally, insects cease feeding behaviours before, during, and after the moult (Ayres & MacLean, 1987; Kang et al., 2019; Rackauskas et al., 2006; Stamp & Bowers, 1994). Similarly, in decapod crustaceans feeding also declines during the pre-moult phase, stops entirely during ecdysis, and resumes only in the late post-moult period (Chan et al., 1988; Dall, 1986; Harpaz et al., 1987). This feeding behaviour reflects the mechanical and physiological challenges posed by exoskeleton shedding, particularly affecting essential feeding structures such as the buccal cavity, oesophagus, and anterior stomach (Lemos & Weissman, 2021). Previous studies indicate that ATP production is regulated by 20E signalling (Chang et al., 2022) and pre-moult is characterized by high energy metabolism which has been supported both at transcriptomic and proteomic level (Head et al., 2019). In various decapod crustaceans, the duration of pre-moult to post-moult can take a substantial amount of time during its moult cycle (Drach & Tchernigovtzeff, 1967; Gorissen & Sandeman, 2022; Kim et al., 2024; Lemos & Weissman, 2021) and the expression of these genes ensures sufficient energy supply during ecdysis.

We also identified genes involved in the melanisation pathway, which plays a key role in cuticle hardening during the post-ecdysial period (Hiruma & Riddiford, 2009). Moreover, we recovered immune-related genes, including components of the Toll and IMD pathways. The melanisation pathway itself occupies an interesting intersection between cuticle maturation and innate immunity, functioning both in cuticle hardening and in pathogen defence (Beckage, 2011; Marmaras et al., 1996; Nakhleh et al., 2017; Vavricka et al., 2010). The vulnerability of arthropods to infection is particularly heightened after ecdysis, when the protective cuticle is incompletely formed. Whether the co-expression of immune genes alongside cuticle maturation reflects the vulnerability that accompanies moulting remains unclear, but previous studies suggest that immune activation may provide protective benefits during this period (Clark & Strand, 2013; Louradour et al., 2017; Tang, 2009; J. Zhang et al., 2014). The melanisation pathway is triggered by PAMPs, such as bacterial peptidoglycan or fungal β-glucans. This activates a protease cascade that converts proPO to active PO, which oxidizes phenols into quinones. These reactive intermediates polymerize to form melanin, which is deposited at sites of infection to encapsulate pathogens and generate cytotoxic reactive oxygen species (ROS) (Beckage, 2011; Nakhleh et al., 2017; Shan et al., 2023; Vavricka et al., 2010). Melanisation has been extensively studied in insects such as *D. melanogaster* and *Bombyx mori*, where it enhances survival during pathogenic infection (Louradour et al., 2017; J. Zhang et al., 2014), interacts with hemolymph coagulation (Bidla et al., 2005), and contributes to broader immune defence (Clark & Strand, 2013; Tang, 2009). Similar immune-related responses during moulting have been observed in decapod crustaceans. In *S. paramamosain*, immune-related genes involved in antimicrobial peptide synthesis, Toll signalling, phenol oxidase system, and antioxidant pathways are significantly upregulated in the hepatopancreas during post-moult and inter-moult stages (Xu et al., 2020). Ortholog co-expression of these gene sets suggests that they play a conserved role during ecdysis.

The co-expression patterns that we observe differ significantly depending on life history strategy. When instar moult stages from holometabolous insect are included, we recover markedly fewer co-expressed genes, particularly those involved in signalling and differentiation pathways. The Hippo and Wnt signalling pathways are important regulators of tissue homeostasis and organ size that coordinate cell proliferation, differentiation and apoptosis (Halder & Johnson, 2011; Komiya & Habas, 2008; Wodarz & Nusse, 1998). In holometabolous insects, growth and differentiation are decoupled (Aldaz & Escudero, 2010; Chapman et al., 2013b; Manthey et al., 2024). During larval development, certain populations of cells, the imaginal primordia, remain in an undifferentiated, pluripotent state with latent embryonic potential and resume differentiation during pupal transition (Aldaz & Escudero, 2010; Chapman et al., 2013b). In holometabolous insects, moulting during the instar stages likely supports mostly growth and cuticle replacement. Extensive studies over the past decades have established that the transition between moults that are dedicated to growth and the extensive differentiation involved in metamorphosis is regulated by a complex interplay of endogenous and environmental factors, involving hormonal feedback between ecdysteroids, *juvenile hormone* (*JH*), and stage-specifying transcription factors such as *Chinmo, Krüppel-homolog 1* (*Kr-h1*), *Broad*, and *E93* (Suzuki et al., 2013; Truman, 2019; Truman et al., 2006; Truman & Riddiford, 2019, 2022). Gradual elaboration of this hormonal crosstalk might have led to the evolution of metamorphosis, by coupling moult cycles to environmental and physiological signals that determine whether an individual proceeds to another juvenile stage or transitions to the adult form. Our results suggest that the moulting programme can be decoupled from broader developmental processes depending on life history strategy.

Surprisingly, the co-expressed genes associated with signalling and differentiation pathways were shared primarily between crustacean moults and insect metamorphic moults rather than with instar larval moults. In insects, EGFR signalling has been implicated in regulating both ecdysteroid and juvenile hormone biosynthesis during developmental transitions (Cruz et al., 2020; Li et al., 2022). Likewise, components of EGFR/ErbB and MAPK signalling are strongly represented in the crustacean Y-organ (Mykles, 2021). Pathways such as EGFR/MAPK signalling might represent a conserved mechanism involved in hormonal regulation of moulting. Although tissue remodelling during crustacean moulting is less extensive than in holometabolous metamorphosis, each moult nevertheless involves coordinated processes of tissue growth and differentiation (Adachi et al., 2025; Covi et al., 2010; Kim et al., 2024). In addition, differentiation pathways such as Wnt signalling have been implicated in setal development in crustaceans (Wang et al., 2023), and setal structures themselves are regenerated during each moult cycle and form the basis of many moult staging systems in decapod crustaceans (Gorissen & Sandeman, 2022; Kim et al., 2024). While the precise functional role of these differentiation pathways in hormonal regulation of moulting in crustaceans remains to be tested, this suggests that metamorphic holometabolous moult might have derived from remodelling in pancrustacean non metamorphic moult.

Despite identifying a small set of genes which are co-expressed across all five species, most of the genes that are differentially expressed during moulting are lineage-specific, either because they do not have orthologs outside of one lineage, or because the differential expression pattern is restricted to one lineage even though the orthologs are more broadly distributed (Figure 3). Temporal co-expression patterns show that these lineage-specific signatures are not evenly distributed across the moulting process. Mid-moulting gene clusters are enriched for relatively younger genes. In *L. vannamei,* these clusters are enriched in functional terms involved in the development of sensory structures, nervous system formation, pigment binding, and chitin-based extracellular matrix components. In contrast, older gene clusters are enriched for functional terms such as steroid metabolic processes, chitin catabolism, and carbohydrate metabolism. This pattern suggests that mid-moult lineage-specific signatures are driven in part by the development of the cuticle and specialized structures of the exoskeleton.

The cuticular surface is remarkably diverse, featuring a wide array of pigmentation and ornamentation, as well as specialized structures (Hallberg & Skog, 2010; Mellon, 2012). This includes various setal structures which line the exoskeleton and are periodically developed with each moult (Drach & Tchernigovtzeff, 1967; Gorissen & Sandeman, 2022; Kim et al., 2024). Many of these are innervated by sensory neurons which may house mechanoreceptive and chemoreceptive neurons, or both (Hallberg & Skog, 2010; Mellon, 2012; Shields et al., 2008). Studies combining physiological analysis with scanning electron microscopy have shown that many cuticular setae on locomotory, feeding, and grooming appendages are bimodal in function, containing both chemo- and mechanoreceptor cells (Hallberg & Skog, 2010; Shields et al., 2008). These structures often serve species-specific roles, including mating behaviour, mechano-sensation, and predator detection. Their diversity and functional specificity across lineages likely explain the recruitment of younger, lineage-specific genes in their regeneration during moulting.

Lineage specific genes have been implicated in species-specific adaptations for behaviour, environmental factors, reproduction, host pathogen interactions, and speciation (Han et al., 2024; Khalturin et al., 2008, 2009). While it is intuitive to conclude that lineage-specific signatures of moulting are due to adaptive divergence, some studies have shown that lineage-specific genes are not always adaptive. Young, lineage-specific genes can be gained, lost, or integrated into gene regulatory networks through stochastic processes, and may persist without signatures of adaptation (Palmieri et al., 2014; Vakirlis et al., 2022; Wacholder et al., 2023). At the same time, the phylogenetic breadth of the species included in this study represents an important limitation. Future studies including additional crustacean lineages, particularly non-malacostracans, as well as other moult types and insect groups such as hemimetabolous insects, will be necessary to determine whether similar lineage-specific gene expression patterns are broadly conserved across Arthropoda. These lineage specific genes thus present interesting candidates for future tests of adaptive evolution in arthropods.

While the overall transcriptome age pattern is similar between *L*. *vannamei* and *D*. *melanogaster*, the underlying gene sets driving these patterns differ substantially. In *L. vannamei*, the mid-moult peak in younger gene expression is primarily driven by genes involved in cuticle development, whereas in *Drosophila*, the same pattern is largely associated with the expression of reproductive genes. Interestingly, in *Drosophila*, gene modules involved with cuticle development are also composed of younger phylostrata, but they are predominantly expressed earlier during pupariation, rather than in mid-moult as in *L. vannamei*. This heterochrony in cuticle gene expression is likely related to the evolution of the puparium in cyclorrhaphous flies, which retain the larval cuticle as a protective case rather than shedding it during ecdysis. Heterochrony in cuticle development and maturation has been linked to adaptation in other arthropods (Elias-Neto et al., 2014; Falcon et al., 2019) which suggests that timing of cuticle formation can evolve in response to different life history and ecological needs.

Although moulting is regulated by conserved hormonal pathways, our findings suggest that it coordinates both deeply conserved and recently evolved gene modules. The precise coordination of these gene modules reflects a regulatory flexibility that enables moulting to accommodate lineage-specific physiological and developmental requirements. It is likely that the regulatory architecture underlying moulting allows for co-expression of genes not directly tied to the moulting process itself but essential for broad physiological support and reconstructing complex structure of the exoskeleton. Since the exoskeleton forms the primary interface between the organism and its environment, environmental pressures may contribute to its morphology and the gene expression programmes involved in its formation. Divergence time and environmental heterogeneity have been shown to influence convergence in both gene expression and morphological traits (Boisseau et al., 2025; Buckley et al., 2009; Chaturvedi et al., 2022). Testing whether ecological niche similarity predicts convergence in ecdysis gene expression would therefore be an interesting task for future work.

By uncovering both conserved and lineage-specific components of moulting gene expression, this study provides evidence for both evolutionary conservation and divergence of this deeply shared developmental process. This conservation of the moulting programme appears to extend to pupariation. These findings suggest that moulting may involve a broader range of developmental processes than its classical definition of periodic shedding of the exoskeleton which may encompass diverse moulting strategies exhibited by Arthropods, ranging from puparium formation to, potentially, apolysis without ecdysis, as observed in paedomorphic insects such as Strepsipterans (Kathirithamby et al., 1984; Manfredini et al., 2007). In a broader perspective, moulting can be viewed as a developmental process that enables growth and morphological transformation through *shedding or reconstruction* of the exoskeleton. Overall, our findings contribute to a broader understanding of the evolutionary dynamics underlying moulting and highlights the modular architecture of the moulting programme, which may be central to understanding how the vast diversity of arthropod species has adapted their post-embryonic development to different ecological and life-history contexts.

## Materials and methods

### Data pre-processing

Table 1 summarizes the RNA sequencing (RNA-seq) data used in this study. Additional time-series datasets were retrieved for *D. melanogaster* from (Ozerova et al., 2025) and for *L. vannamei* from Gao et al., 2015b (Supplementary Table 1). The quality of the reads was checked using FastQC v.0.11.2 (Andrews 2010) and adapter trimming and quality control were performed using Trimmomatic v. 0.36 (Bolger et al., 2014) (Supplementary Table 5). Trimmed reads were pseudo-aligned against the respective transcriptome of each species using kallisto v0.44.0 (Bray et al., 2016). Transcriptome and annotation files were retrieved from National Center for Biotechnology Information (NCBI) databases (Sayers et al., 2024). The quality and completeness of the genome assembly of each species were assessed with Benchmarking Universal Single-Copy Orthologs (BUSCO) using the Arthropoda lineage dataset (odb10) as reference (Simão et al., 2015) through the A3CAT database (Feron & Waterhouse, 2022). All assemblies used in this study have a BUSCO score of more than 90% (Supplementary Table 6). Transcript level expression was imported into R v4.2.2 and summarised to the gene level using the R/tximport v1.10.1 (Soneson et al., 2016).

### Construction of orthogroup gene expression matrix

To examine gene expression patterns of moulting stages between groups of species, orthologous genes (orthogroups) were identified using Orthologer v3.0.2 with default parameters (Kuznetsov et al., 2023) after filtering the proteomes for the single longest protein-coding isoform of each gene. Orthology inference was performed using a broader dataset including additional crustacean species (*Penaeus japonicus* and *Eriocheir sinensis*) beyond the five species in this study in order to improve orthology assignment associated with limited taxonomic sampling. The cross-species expression matrix was constructed from orthogroups shared across all five species and was not restricted to strict one-to-one orthologs. Because some orthogroups contained multiple genes per species, a single representative gene per species was selected based on the highest mean expression across samples, generating a read count matrix of 5,543 orthogroups identified across the five species. This orthogroup matrix was used exclusively for PCA and clustering analysis, whereas differential expression analyses were performed independently within each species using the original gene-level datasets. To assess the robustness of the representative gene selection strategy, we also used an alternative orthogroup matrix in which a random representative gene was selected per orthogroup. This produced similar overall PCA patterns, indicating that the major biological signal is robust to the choice of orthogroup representative (Supplementary Figure S6).

### Principal component analysis

Prior to conducting cross-species principal component analysis (PCA), PCA was first performed separately for each species to assess sample clustering of moulting stages within each dataset. Combined raw read counts from the 5,543 orthogroups identified across the five species were transformed using variance stabilizing transformation (VST function) implemented in DESeq2 (Love et al., 2014). This transformation generated constant variances across all read counts, creating a homoscedastic dataset. To account for the effect of species on gene expression, we included species as an explanatory variable in the model when constructing the read counts matrix. A PCA was conducted to examine the overall gene expression pattern of the dataset using the PCAtools R package (Blighe and Lun 2022) on the VST RNA-seq counts. The top 10 PCs were selected, and individual eigenvalues were tested for correlation with the traits of interest: taxonomic group, moulting stage, and species.

### Differential gene expression analysis

Differential gene expression analyses between pre-moult and post-moult stages were performed individually per species using DESeq2 (Love et al., 2014), with P-values adjusted using the Benjamini–Hochberg (Benjamini & Hochberg, 1995) procedure to control for false discovery rate (FDR). An adjusted *P*-value of 0.05 was used as the threshold for identifying differentially expressed genes (DEGs). For simplicity, wandering and prepupal stages of the larval-prepupal moult are referred to as simply pre-moult and post-moult, respectively. The resulting differentially expressed genes from each species were grouped based on orthology of DEGs into the following categories: species specific genes (differential expression not shared with any other species), Pancrustacea shared genes (differential expression in at least one insect and one malacostracan), class specific genes restricted to either class Insecta or Malacostraca (differential expression in at least two species per group), and genes which have differential expression exclusive to metamorphic or to non-metamorphic samples. For each DEG, we also assessed whether its corresponding orthologous group, irrespective of differential expression, was restricted to a single lineage (insect-only or malacostracan-only), shared between insects and malacostracans (Pancrustacea-shared), or lacked detectable orthologs in other taxa. We note that genes classified as having no detectable orthologs may nevertheless have orthologs in additional closely related taxa not included in the current comparison, such as other Dipterans. Thus, a species-specific DEG may either represent a gene with no detectable ortholog in the present dataset, or the only differentially expressed member of an orthogroup otherwise shared across multiple species in the study.

### Clustering and functional analysis

Clust (Abu-Jamous & Kelly, 2018) was used to identify and extract distinct groups of genes displaying correlated expression patterns across all samples. Unlike standard clustering approaches, Clust does not require pre-specifying a fixed number of clusters. Briefly, Clust applies k-means clustering using a range of k-values to generate an initial set of seed clusters, which are subsequently refined using M–N scatter plots to identify large, non-overlapping clusters with low dispersion. As such, the final number of clusters is determined dynamically by the data rather than fixed a priori. We used the default of Clust which filters out genes with flat expression profiles prior to analysis. Several cluster tightness values ranging from 0.1 to 1 were evaluated. Smaller tightness values produce larger and more diffuse clusters, whereas larger values produce smaller and more tightly co-expressed clusters. A tightness value of 1 was selected for this study as it produced clusters with minimal dispersion and clear expression patterns. Gene Ontology enrichment analysis (Ashburner et al., 2000) of genes belonging to each cluster was performed using ClusterProfiler (Wu et al., 2021). Putative cuticular protein families were identified using CutProtFam-Pred (Ioannidou et al., 2014), which uses Hidden Markov Models based on CuticleDB (Magkrioti et al., 2004) to detect and classify cuticular protein families based on conserved sequence motifs, including cuticular protein subfamilies Rebers and Riddiford type 1 (RR-1) and type 2 (RR-2), as well as additional families such as tweedle and cuticular protein analogous to peritrophins types 1 and 3 (CPAP1 and CPAP3).

### Transcriptome age analysis and Phylostratigraphy

To examine the conservation of gene expression across different timepoints of moulting, phylostratigraphy analysis was used to estimate the relative transcriptome age of the time-series samples (Supplementary Table 1) (Domazet-Lošo & Tautz, 2010). VST-transformed counts were generated using DESeq2 (Love et al., 2014), and mean expression values for each stage were calculated. The resulting gene expression matrix was then clustered using Clust (Abu-Jamous & Kelly, 2018), and the phylostratum composition for each cluster was determined by using GenEra v1.4.0 (Barrera-Redondo et al., 2023). All genes of a given target genome were queried against a protein database built from eukaryotic reference genomes retrieved from NCBI using DIAMOND *blastp* in more-sensitive mode with an *e*-value threshold of 10⁻⁵ (Buchfink et al., 2015). *Blastp* hits were considered homologues of the query protein and were used to determine the phylostrata of the gene.

The Transcriptome Age Index (TAI) of RNA-seq samples was calculated as described previously (Domazet-Lošo & Tautz, 2010) using the *myTAI* (v2.0.0) (Drost et al., 2018) package in R (v4.3.2). TAI values were computed as the weighted mean of the phylostratum rank of a given gene based on its expression level in the transcriptome of an RNA-seq sample. VST transformed gene expression was used, which like log-transform removes the impact of extreme values (see J. Liu & Robinson-Rechavi, 2018; Piasecka et al., 2013). A high TAI value indicates a younger transcriptome age, whereas a lower TAI corresponds to an older transcriptome. Statistical significance of the TAI profile was determined through a permutation-based test implemented in *myTAI* (v2.0.0) (Drost et al., 2018). This test assesses whether the overall TAI trajectory deviates from a random expectation and therefore evaluates the global structure of transcriptome age dynamics across the moulting process rather than pairwise differences between individual stages.

## Data availability

All data used in this study were publicly available datasets retrieved from NCBI with the following accession numbers. *D. melanogaster* (PRJNA75285, PRJNA1124160), *Z. cucurbitae* (PRJEB36344, PRJNA562032), *B. dorsalis* (PRJNA524281), *L. vannamei* (PRJNA288849), *S. paramamosain* (PRJNA687923). Additional supplementary files and scripts used in this study are available at Zenodo: *10.5281/zenodo.17784461*.

## Acknowledgments

The authors would like to thank all members of the SNSF Sinergia Arthropod Moulting Project: Michele Leone, Valentine Rech de Laval, Ariel Chipman, Olga Volovych, Idan Sheizaf, Asya Novikova, Sinead Lynch, Jonathan Antcliffe, Harriet Drage and Allison Daley, and the Robinson-Rechavi group and Christabel Bucao (EMBL), for their invaluable support, and insightful discussions throughout this study. This work was supported by funding from a Swiss National Science Foundation (SNSF) Sinergia award [grant number 198691]. Silhouette icons used in this study were all retrieved from Phylopic under Universal Public Domain License (CC0 1.0) and Public Domain Mark (PDM).

