## Supplementary Figures for "Moulting in Pancrustacea is characterised by both deeply conserved and recently evolved gene modules"

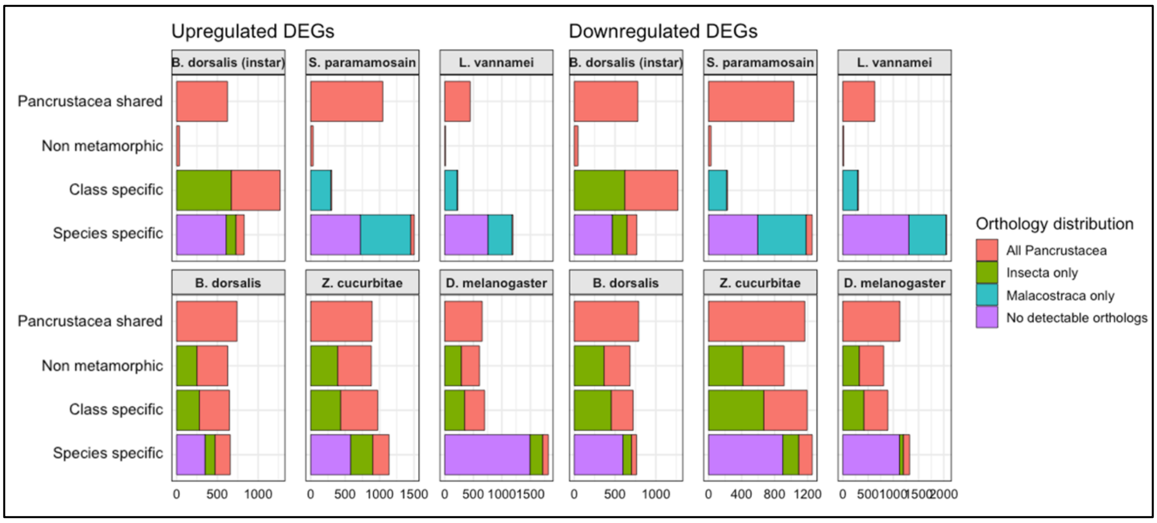


**Figure S1.** Barplot showing summary of differentially expressed genes (DEGs) per species grouped by different categories. Left panel shows upregulated DEGs per species. Right panel shows downregulated DEGs per species.

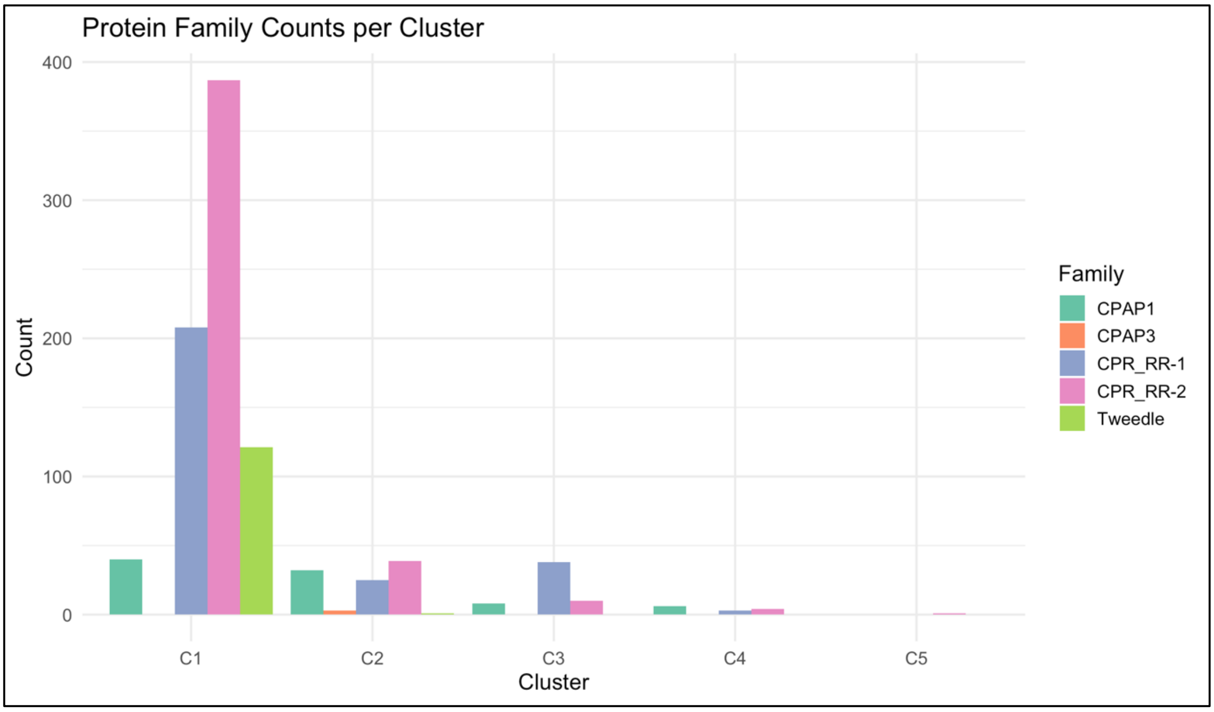


**Figure S2.** Barplot showing total number of cuticle protein families in *L. vannamei* detected per cluster

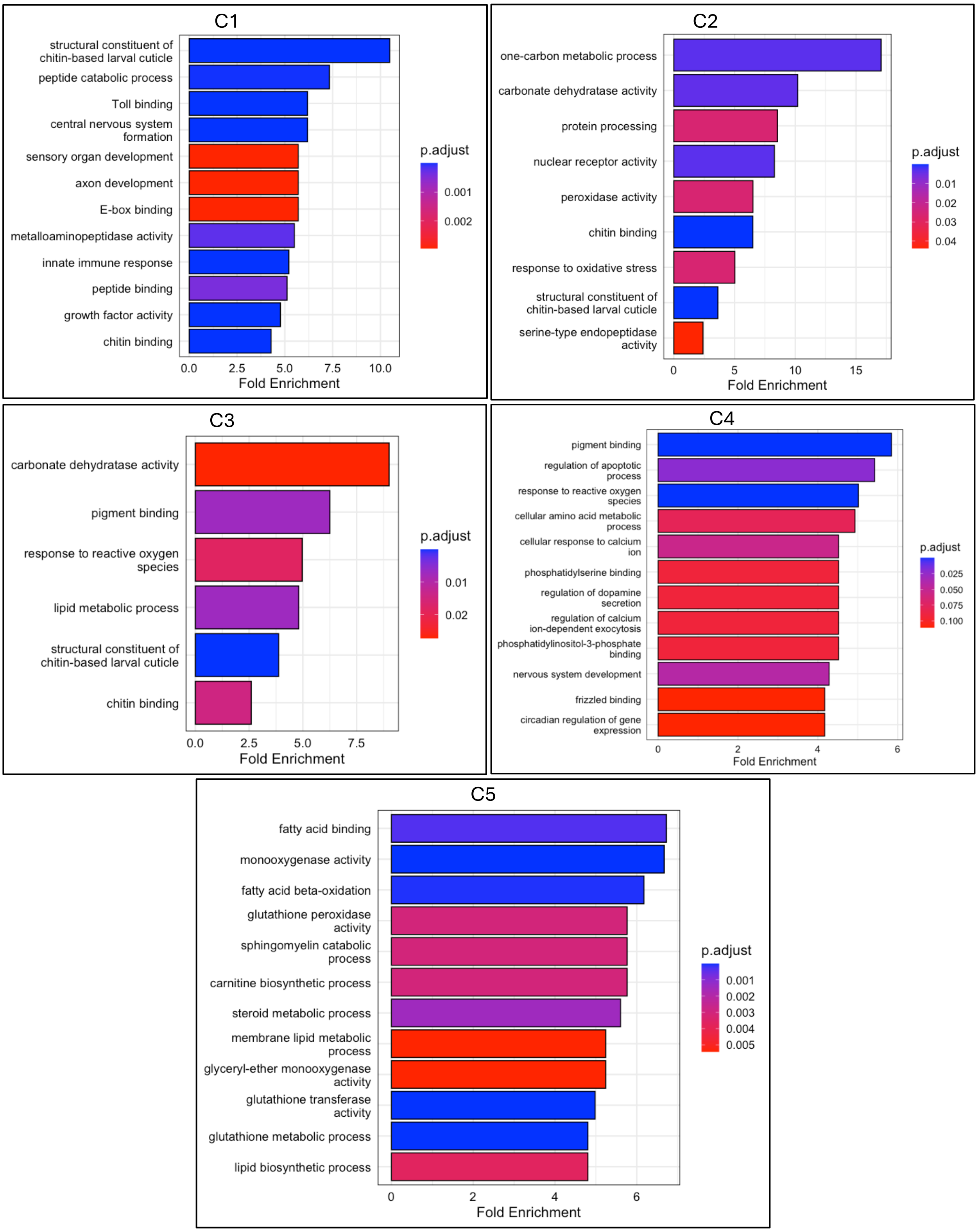


**Figure S3.** GO enrichment plots showing significantly enriched terms per gene expression cluster in *L. vannamei*

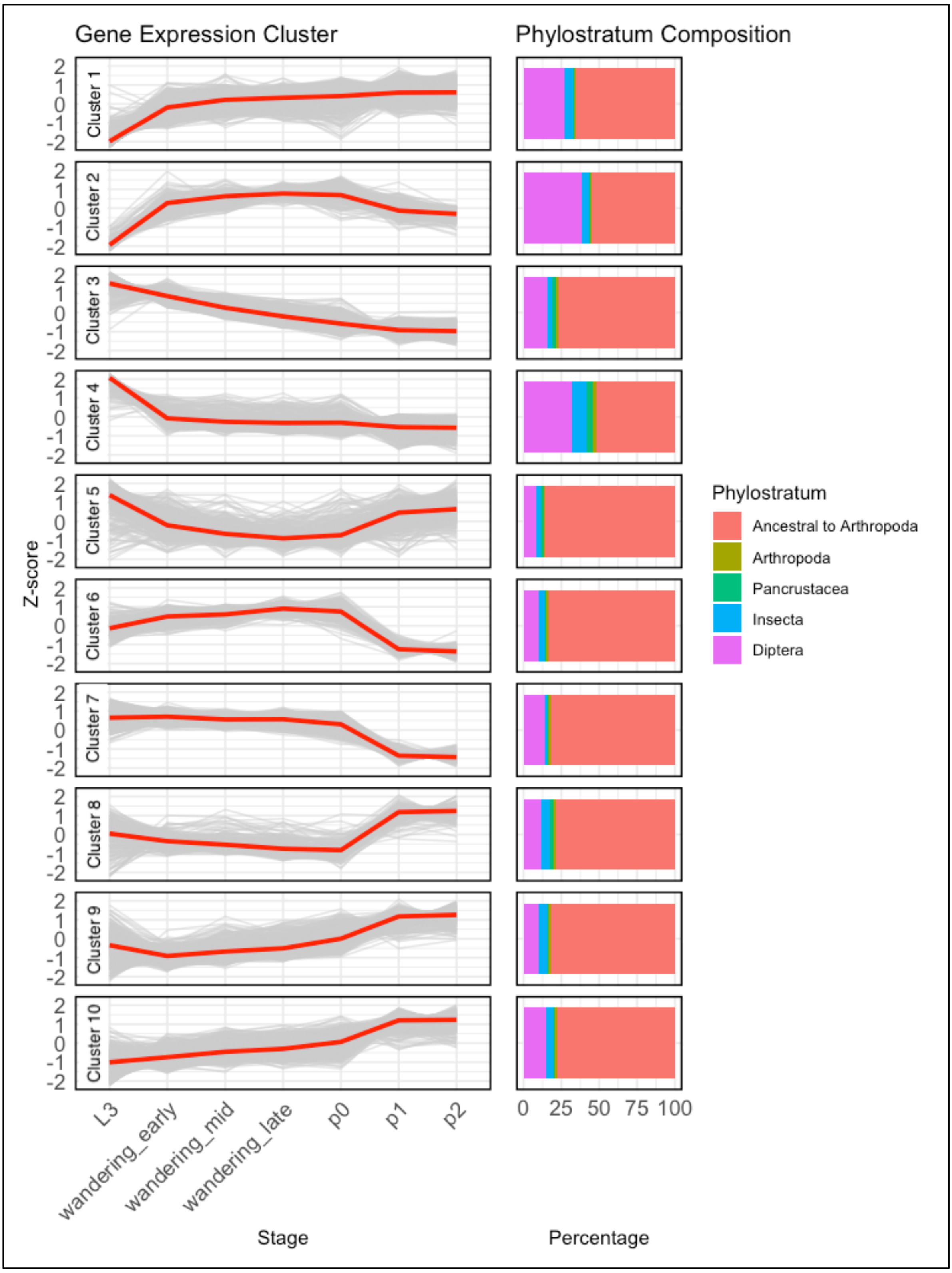


**Figure S4. Moulting gene expression clusters in *D. melanogaster***
Figure above shows the gene expression clusters of *D. melanogaster* samples and their corresponding phylostratum composition.

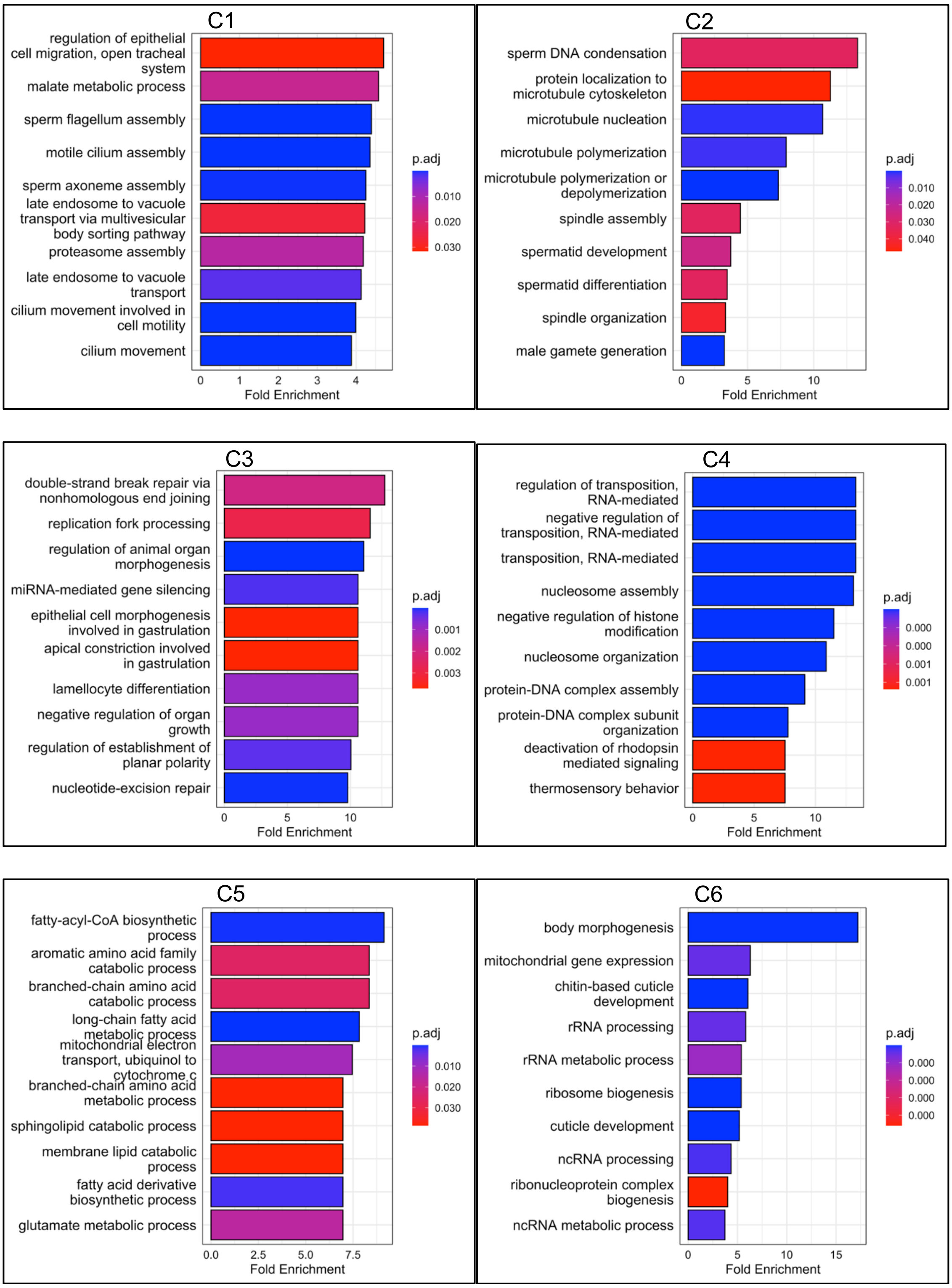
C1
regulation of epithelial cell migration, open tracheal system
malate metabolic process
sperm flagellum assembly
motile cilium assembly
sperm axoneme assembly
late endosome to vacuole transport via multivesicular body sorting pathway
proteasome assembly
late endosome to vacuole transport
cilium movement involved in cell motility
cilium movement
0 1 2 3 4
Fold Enrichment
p.adj
0.010
0.020
0.030
C2
sperm DNA condensation
protein localization to microtubule cytoskeleton
microtubule nucleation
microtubule polymerization
microtubule polymerization or depolymerization
spindle assembly
spermatid development
spermatid differentiation
spindle organization
male gamete generation
0 5 10
Fold Enrichment
p.adj
0.010
0.020
0.030
0.040
C3
double-strand break repair via nonhomologous end joining
replication fork processing
regulation of animal organ morphogenesis
miRNA-mediated gene silencing
epithelial cell morphogenesis involved in gastrulation
apical constriction involved in gastrulation
lamellocyte differentiation
negative regulation of organ growth
regulation of establishment of planar polarity
nucleotide-excision repair
0 5 10
Fold Enrichment
p.adj
0.001
0.002
0.003
C4
regulation of transposition, RNA-mediated
negative regulation of transposition, RNA-mediated
transposition, RNA-mediated
nucleosome assembly
negative regulation of histone modification
nucleosome organization
protein-DNA complex assembly
protein-DNA complex subunit organization
deactivation of rhodopsin mediated signaling
thermosensory behavior
0 5 10
Fold Enrichment
p.adj
0.000
0.000
0.001
0.001
C5
fatty-acyl-CoA biosynthetic process
aromatic amino acid family catabolic process
branched-chain amino acid catabolic process
long-chain fatty acid metabolic process
mitochondrial electron transport, ubiquinol to cytochrome c
branched-chain amino acid metabolic process
sphingolipid catabolic process
membrane lipid catabolic process
fatty acid derivative biosynthetic process
glutamate metabolic process
0.0 2.5 5.0 7.5
Fold Enrichment
p.adj
0.010
0.020
0.030
C6
body morphogenesis
mitochondrial gene expression
chitin-based cuticle development
rRNA processing
rRNA metabolic process
ribosome biogenesis
cuticle development
ncRNA processing
ribonucleoprotein complex biogenesis
ncRNA metabolic process
0 5 10 15
Fold Enrichment
p.adj
0.000
0.000
0.000
0.000

(cont.)

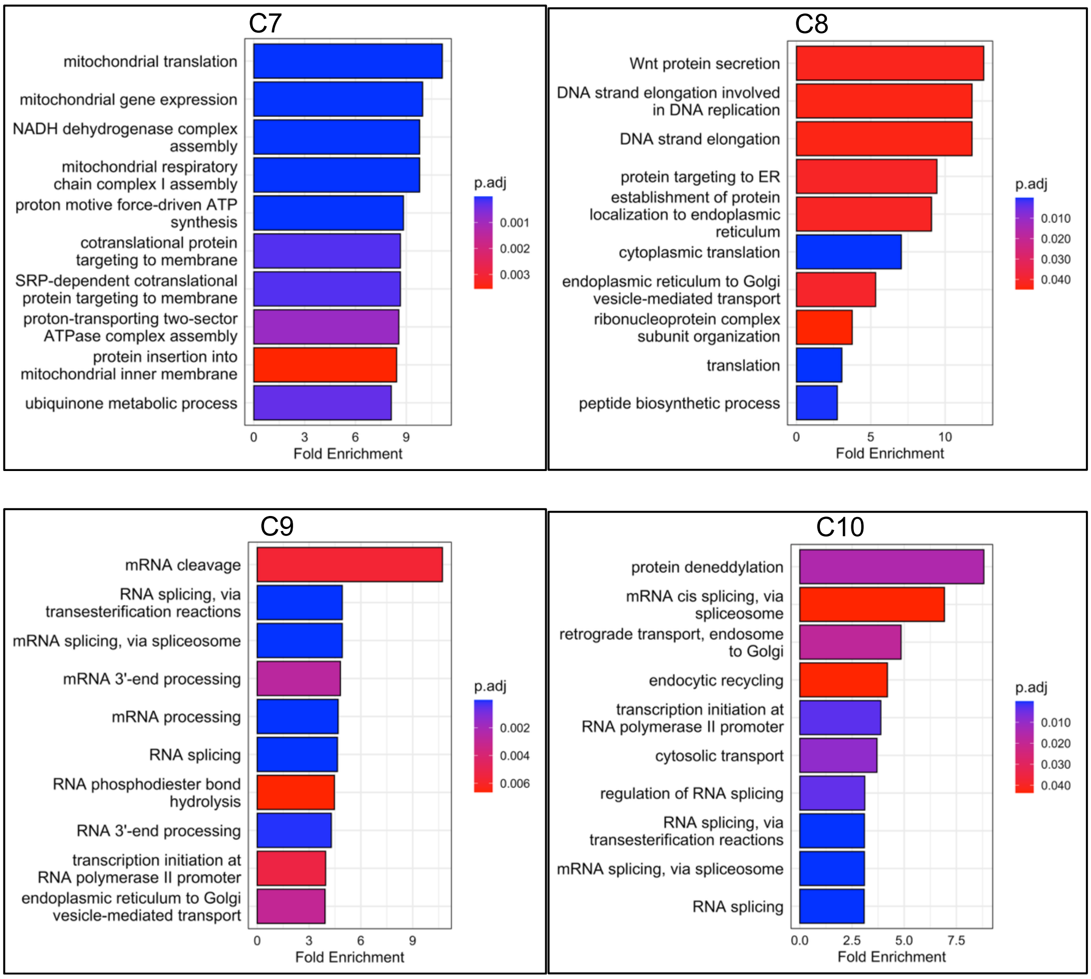


**Figure S5.** GO enrichment plots showing significantly enriched terms per gene expression cluster in *D. melanogaster*

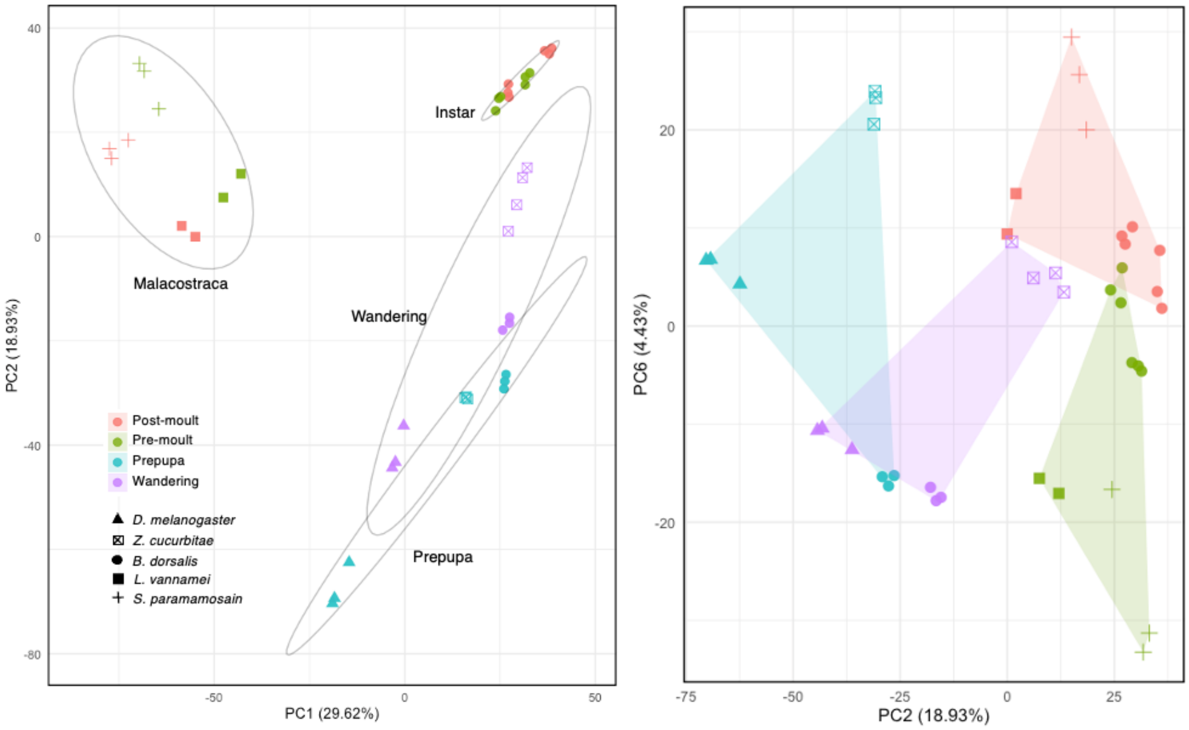


**Figure S6.** PCA using an alternative orthogroup matrix in which a random representative gene was selected per orthogroup. Left plot showing the PC1 and PC2 and right plot showing PC2 and PC6.

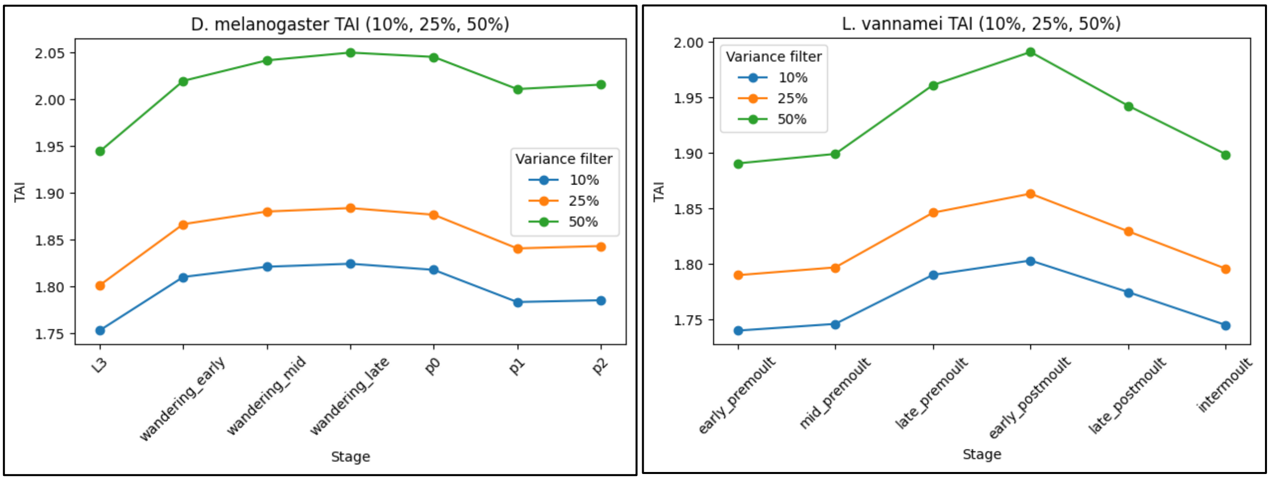


**Figure S7.** TAI of various moult stages in *D. melanogaster* and *L. vannamei* filtered based on expression variance.
